# A metaproteomics method to determine carbon sources and assimilation pathways of species in microbial communities

**DOI:** 10.1101/245290

**Authors:** Manuel Kleiner, Xiaoli Dong, Tjorven Hinzke, Juliane Wippler, Erin Thorson, Bernhard Mayer, Marc Strous

## Abstract

Measurements of the carbon stable isotope ratio (δ^13^C) are widely used in biology to address major questions regarding food sources and metabolic pathways used by organisms. Measurement of these so called stable carbon isotope fingerprints (SIFs) for microbes involved in biogeochemical cycling and microbiota of plants and animals have led to major discoveries in environmental microbiology. Currently, obtaining SIFs for microbial communities is challenging as the available methods either only provide limited taxonomic resolution, such as with the use of lipid biomarkers, or are limited in throughput, such as NanoSIMS imaging of single cells.

Here we present “direct Protein-SIF” and the Calis-p software package (https://sourceforge.net/projects/calis-p/), which enable high-throughput measurements of accurate δ^13^C values for individual species within a microbial community. We benchmark the method using 20 pure culture microorganisms and show that the method reproducibly provides SIF values consistent with gold standard bulk measurements performed with an isotope ratio mass spectrometer. Using mock community samples, we show that SIF values can also be obtained for individual species within a microbial community. Finally, a case study of an obligate bacteria-animal symbiosis showed that direct Protein-SIF confirms previous physiological hypotheses and can provide unexpected new insights into the symbionts’ metabolism. This confirms the usefulness of this new approach to accurately determine δ^13^C values for different species in microbial community samples.

**Significance:** To understand the roles that microorganisms play in diverse environments such as the open ocean and the human intestinal tract, we need an understanding of their metabolism and physiology. A variety of methods such as metagenomics and metaproteomics exist to assess the metabolism of environmental microorganisms based on gene content and gene expression. These methods often only provide indirect evidence for which substrates are used by a microorganism in a community. The direct Protein-SIF method that we developed allows linking microbial species in communities to the environmental carbon sources they consume by determining their stable carbon isotope signature. Direct Protein-SIF also allows assessing which carbon fixation pathway is used by autotrophic microorganisms that directly assimilate CO_2_.

## Introduction

Measurements of stable carbon isotope ratios (^13^C/^12^C) are used in many different scientific fields, including atmospheric sciences, biology, paleoclimatology, oceanography, geology, environmental sciences, food and drug authentication, and forensic applications (1). In biology, stable isotope ratios (Stable Isotope Fingerprints = SIF) can be used to address at least two major questions. First, what is the food source of an organism? This question can be answered based on the principle that heterotrophic organisms usually have a similar SIF to their food source (“you are what you eat”) (2). This has been used, for example, to assess the diet of animals (3) and to determine which microorganisms consume a specific carbon source (e.g. methane) in marine sediments (4). The second question, which can be addressed for those organisms that grow on C_1_ carbon sources (bicarbonate, CO_2_, methane), is which metabolic pathway is used to assimilate the carbon source? This question can be answered based on the principle that most metabolic pathways/enzymes for C_1_ assimilation discriminate against ^13^C, which leads to characteristic carbon isotope fractionation effects. The extent of isotope fractionation of different C_1_ assimilation pathways varies and thus the metabolic pathway used can be predicted based on the extent of the isotope fractionation (2, 5). This has, for example, been used in the past to distinguish plants with different carbon assimilation physiologies (6) and to predict differences in carbon fixation pathways used by symbionts of marine animals (7, 8).

Obtaining the SIFs of individual species in microbial communities is in theory a very promising tool to help unravel important abiotic and biotic interactions in global biogeochemical cycles as well as in microbiota of plants and animals. For example, if SIFs of individual species in the intestinal microbiota of humans were known, we could deduce which dietary components are used by different species in the intestine. However, there is currently no experimental approach to determine the specific SIFs of a large number of species in communities with reasonable effort and cost. The presently available approaches either have no or limited taxonomic resolution or are low throughput. The most common approach for measuring ^13^C/^12^C ratios, isotope ratio mass spectrometry (IRMS), usually determines highly accurate C isotope ratios for bulk organic samples that have been converted to the measurement gas CO2 via thermal decomposition. In IRMS the measured C isotope ratios are reported as δ^13^C values, which give the per mille (‰) deviation of the measured ratio from the internationally accepted standard V-PDB (Vienna Peedee Belemnite). If lipid biomarkers are separated followed by their C isotope analysis using IRMS, high level taxonomic groups can be resolved (9). Recently separation of proteins has also been combined with C isotope analysis using IRMS (P-SIF) (10). The P-SIF approach theoretically allows assigning δ^13^C values to 5-10 taxa per sample, however, the approach has only been used on two bacterial pure cultures so far as it is extremely low throughput because the mass spectrometer run time required for peptide identification alone amounts to around two weeks per sample. A final approach is the combination of fluorescence *in situ* hybridization and nanoSIMS, which enables measurement of δ^13^C values of individual cells (4), however, this approach has a low throughput because specific fluorescently labeled probes have to be applied for each individual species or higher level taxonomic group.

Here we present “direct Protein-SIF” and the Calis-p (The CALgary approach to ISotopes in Proteomics) software package, a new method that enables high-throughput measurement of δ^13^C (SIF) values for individual species within microbiota and environmental microbial communities using metaproteomics. We use the word “direct” to highlight the fact that the SIF data is directly extracted from a standard metaproteomic dataset, i.e. the same mass spectrometry data is used for both peptide identification and SIF estimation. The direct Protein-SIF workflow (Fig. 1 and S1) consists of a standard proteomics sample preparation to produce peptide mixtures of the samples and a reference material, followed by acquisition of 1D- or 2D-LC-MS/MS data of the peptide mixture using a high-resolution Orbitrap mass spectrometer (for details see the methods section). The MS^2^ data is used as input for a standard proteomic database search to produce a table with scored peptide-spectral matches (PSMs). The PSM tables for the samples and the reference material plus the raw mass spectrometry data (in mzML format) are used as input for the Calis-p software. In a first step the software finds the isotopic peaks for each PSM and sums their intensity across a retention time window. The isotope peak intensities are reported together with the sum formula of the identified peptides. Isotope peak intensity patterns with low intensities or low search engine scores are discarded. The remaining isotope patterns are used as input for the stable isotope fingerprinting. In this step, experimentally derived isotope peak distributions are fitted to theoretical isotope peak distributions computed for a range of δ^13^C values with a Fast Fourier Transform method adapted from Rockwood et al. (11). Peptides with a poor goodness of fit are discarded. The remaining peptides are used to compute an average δ^13^C value for each individual species and associated standard errors in ‰. In a final step the species δ^13^C values are corrected for instrument isotope fractionation by applying the offset determined using the reference material (see details in the results section).

**Figure 1:**
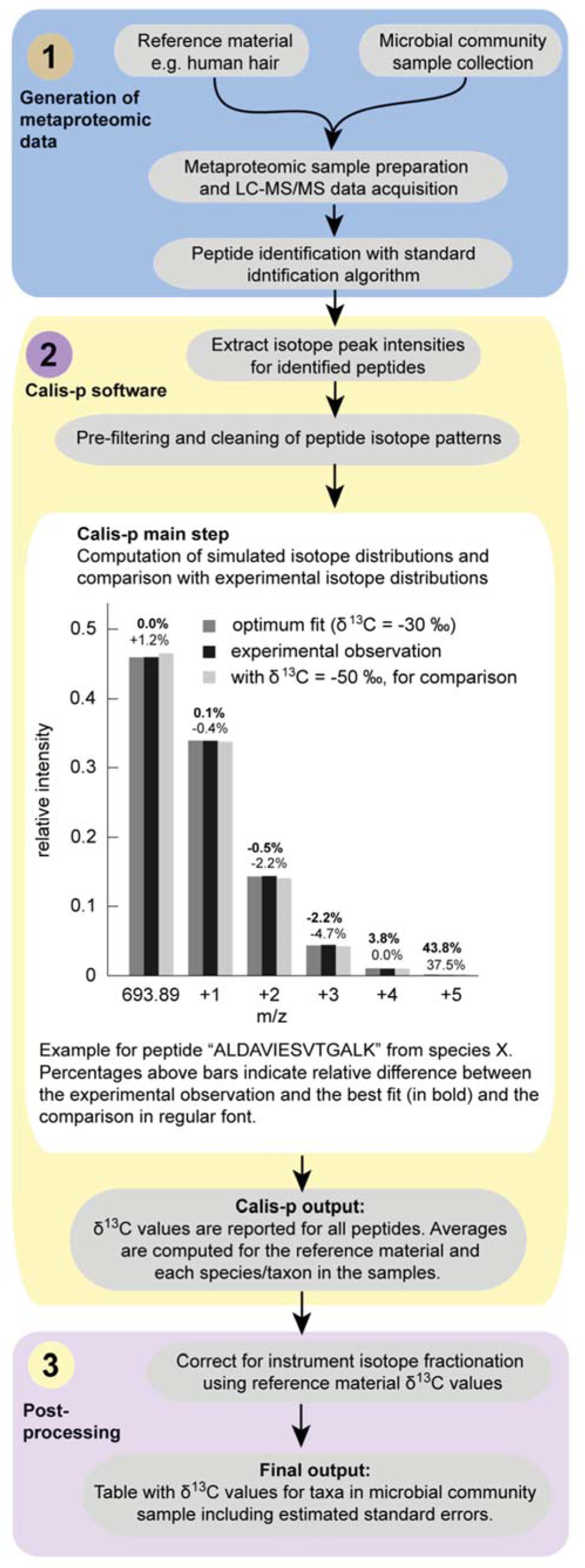
Direct Protein-SIF workflow. In the main step of the data analysis with the Calis-p software, the experimentally derived isotope distributions for peptides are compared with theoretical isotope peak distributions computed based on peptide molecular formulae with a Fast Fourier Transform method. The comparison with theoretical distributions is done for a specified range of δ^13^C values in increments. Goodness of fit is calculated for all comparisons and the δ^13^C value for the best fit is reported if a pre-determined goodness of fit threshold is passed. A more detailed workflow can be found in Figure S1.

## Results

### Benchmarking with pure culture data

For benchmarking, we measured the stable carbon isotope ratios of 20 pure culture species using both direct Protein-SIF and Continuous Flow-Elemental Analysis-Isotope Ratio Mass Spectrometry (CF-EA-IRMS). CF-EA-IRMS is the most commonly used method to determine highly accurate carbon isotopic compositions of organic bulk samples with measurement uncertainties of ± 0.15 ‰ or less and can be considered the gold standard. The 20 pure cultures represented 18 bacterial, one archaeal, and one eukaryotic species (Tables S1 and S2). For seven of the species we obtained technical replicate measurements for the direct Protein-SIF method.

The Protein-SIF δ^13^C values for the technical replicate measurements of individual species were highly consistent with each other and generally deviated by less than ±1‰ from the mean (Table S1). The direct Protein-SIF δ^13^C values were linearly correlated to the CF-EA-IRMS values (R^2^ = 0.94). The direct Protein-SIF δ^13^C values showed a systematic offset from the CF-EA-IRMS values of -15.4‰ (SD = 2.55) (Fig. 2a). This systematic offset is likely caused by isotope fractionation in the Orbitrap mass spectrometer, which has been recently described (12). The large range of δ^13^C values measured with both direct Protein-SIF and CF-EA-IRMS corresponds to the isotope ratios of the different carbon sources used to grow the pure cultures (Table S3) and the heterotrophic versus autotrophic lifestyle of the different microorganisms.

**Figure 2:**
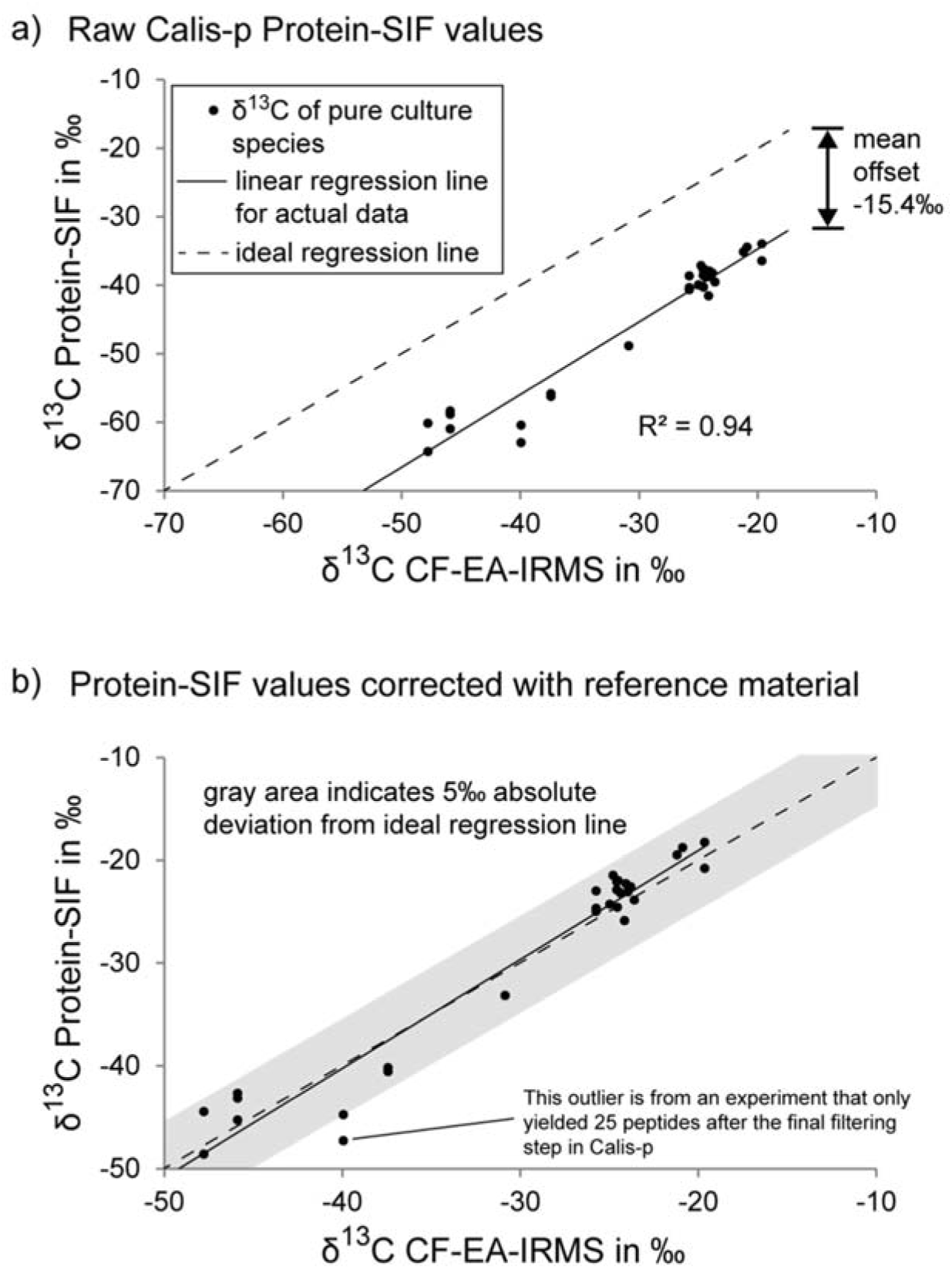
Comparison of δ^13^C measurements of pure cultures with Protein-SIF and CF-EA-IRMS. Twenty pure cultures representing 18 bacterial, one archaeal and one eukaryotic species were measured with both methods (detailed data in Tables S1 and S2). For seven of the species technical replicate measurements were obtained. a) The raw δ^13^C values from the Calis-p software plotted against the IRMS derived values. The average offset of Protein-SIF values from the IRMS values is indicated. b) Protein-SIF values after offset correction using the offset determined with reference material (human hair).

### Correction of instrument isotope fractionation with reference material

To correct for instrument isotope fractionation when using the direct Protein-SIF method we implemented the use of a reference material which is prepared, measured and analyzed alongside the samples. With CF-EA-IRMS, reference materials with known isotopic compositions are also needed to correct for instrument isotope fractionation (13). We chose human hair as the reference material for the following reasons: (i) The bulk of its dry mass consists of protein, which allows to directly correlate Protein-SIF and IRMS δ^13^C values. Interference from other biomolecules with potentially different isotope composition, such as lipids, is limited. (ii) It contains a large diversity of proteins providing hundreds to thousands of different peptides that can be measured for Protein-SIF. (iii) The protein sequences required for peptide identification are known. (iv) It is easy to obtain in large batches (a few grams) to serve as a reference material for many years. Human hair with known δ^13^C values measured by IRMS is also available from major chemical suppliers and the U.S. Geological Survey (https://isotopes.usgs.gov/lab/referencematerials.html).

The offset between δ^13^C values generated with direct Protein-SIF and CF-EA-IRMS for the human hair reference material was -15.7‰ and thus almost identical to the average offset for the benchmarking pure culture δ^13^C values (Fig. 2a). We corrected the pure culture Protein-SIF δ^13^C values with the reference material offset. After this, the absolute deviation of the Protein-SIF δ^13^C values from the CF-EA-IRMS δ^13^C values was on average ±2.1‰ (SD=1.5) (Fig. 2b, Table S2).

### Detection limit and accuracy of Protein-SIF in mock communities

To determine the detection limit of our approach and to test how accurately we can detect SIFs in complex community samples, we used mock communities. These mock communities were created by mixing the 20 pure culture species used for benchmarking with 12 additional microbial strains and species so that the final community contained a total of 32 strains and species. We generated a total of 12 biological replicates of the community with differing species abundances and analyzed each replicate with two different LC-MS/MS methods (14). To evaluate the mock community direct Protein-SIF data we only considered the 20 benchmarking species in the community for which IRMS and direct Protein-SIF δ^13^C values were known based on the results presented above (“Benchmarking with pure culture data”).

The accuracy and detection limit of direct Protein-SIF depends heavily on the available amount of data. We found that the number of peptides for each species that pass the final Calis-p filtering step has a large influence on the accuracy of the determined SIF values (Fig. 3). If less than 60 peptides were available for SIF value calculation, the Protein-SIF δ^13^C values differed from the IRMS derived δ^13^C values by more than 10‰ for more than half of the 20 species. If between 60 and 100 peptides were available, protein-SIF values differed less than 10‰ from IRMS values for 6 out of 9 species, with only one large outlier. With more than 100 peptides available, over 95% of Protein-SIF values differed less than 10‰ from δ^13^C values determined by IRMS (nean deviation 4.7‰). As expected, the detection limit for direct Protein-SIF depended on the amount of mass spectrometric data available, because to reach the necessary number of high quality peptide isotopic patterns for low abundant species more mass spectra are required (Table 1 and S4). For example, one four-hour LC-MS/MS run enabled determination of SIF values for 25% of the species in the mock community. These species were relatively abundant (proteinaceous biomass > 5.7% of the total community). In contrast, combined data of twelve four-hour runs enabled determination of SIF values for 75% of species, including species with lower (< 1% of the total proteinaceous biomass) abundance (Table 1).

**Figure 3:**
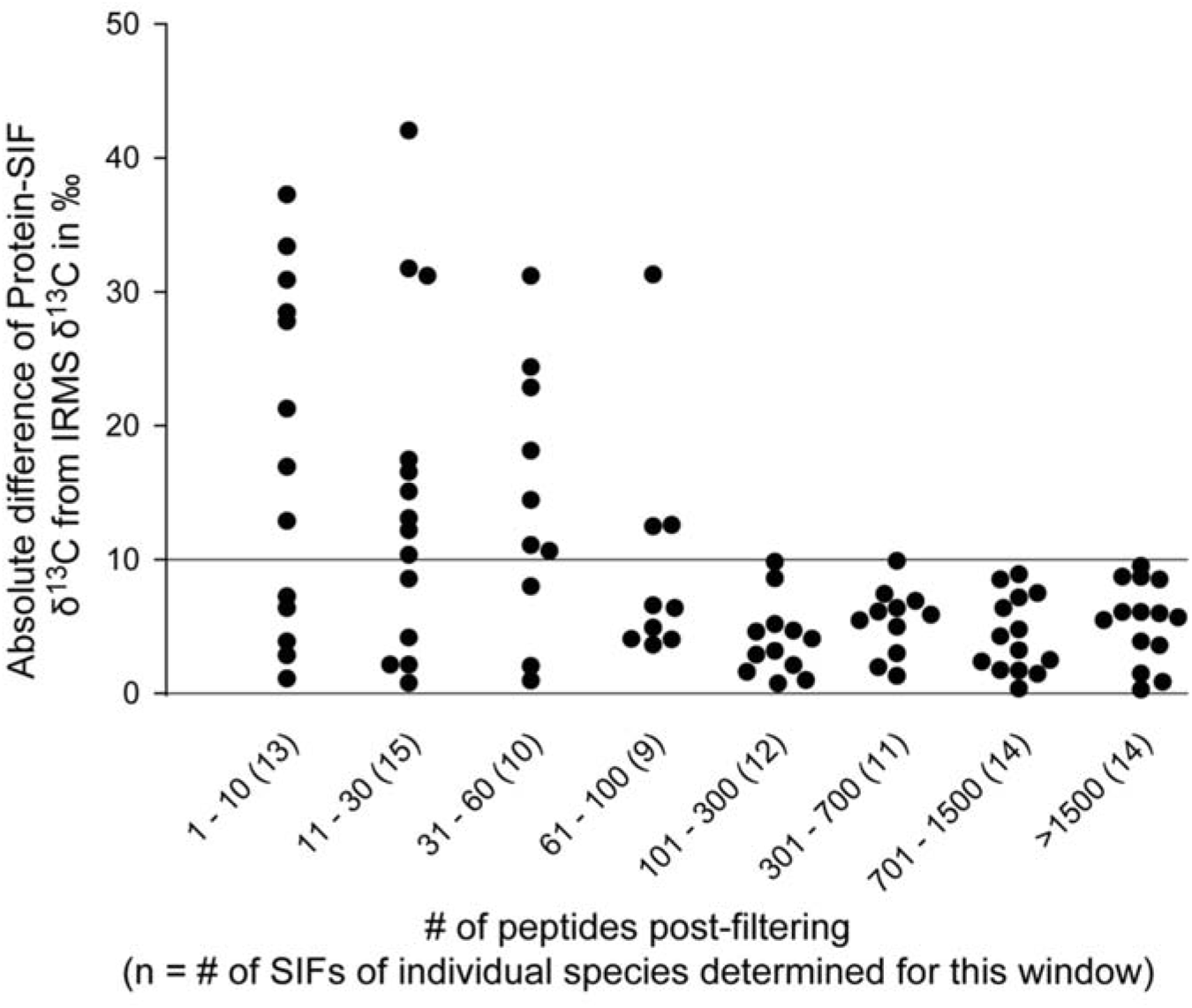
Absolute difference between δ^13^C values determined with direct Protein-SIF of individual species in mock communities and IRMS of the corresponding pure cultures. Five mock community datasets with a total of 32 species and strains were analyzed. For 20 species, the δ^13^C values were known from IRMS performed on pure cultures. For these species, the δ^13^C values were determined with direct Protein-SIF. Each dataset contained different amounts of data (see Table 1). Different numbers of peptides were identified and passed the final Calis-p peptide filter for each species in each dataset. The absolute difference between δ^13^C values obtained via Protein-SIF and IRMS was calculated and sorted according to how many peptides were available for SIF calculation by Calis-p after filtering the peptides. The plot gives the absolute differences for different ranges of peptide # used for SIF calculation.

**Table 1:** Detection limit of direct Protein-SIF for species in mock communities depending on the amount of LC-MS/MS data available^1^.

| total LC-MS/MS run time | 92 hours | 52 hours | 31 hours | 8 hours | 4 hours |
| --- | --- | --- | --- | --- | --- |
| Biological replicates | 12 | 12 | 4 | 1 | 1 |
| Gradient length (min) | 460 | 260 | 460 | 460 | 260 |
| MS<sup>2</sup> spectra (in millions) | ~2.04 | ~1.2 | ~0.68 | ~0.17 | ~0.1 |
| Species (out of 20) with Protein-SIF<sup>2</sup> | 15 | 15 | 10 | 6 | 5 |
| Lower species abundance (%) limit for SIF determination<sup>3</sup> | 0.82 | 0.82 | 0.92 | 5.65 | 5.79 |
| Mean deviation of Protein-SIF $\delta^{13}\text{C}$ from IRMS $\delta^{13}\text{C}$ in ‰ | 3.6 | 5.4 | 4.7 | 4.4 | 5.8 |
| Min deviation in ‰ | 0.2 | 0.6 | 0.9 | 1.5 | 2.8 |
| Max deviation in ‰ | 9.4 | 9.7 | 8.8 | 8.4 | 9.8 |

2: Number of species with a sufficient number of peptides for Protein-SIF (>100) after final Calis-p filtering.
3: Abundance (% of total community protein) of lowest abundance species for which determining Protein-SIF value was possible i.e. >100 peptides after final Calis-p filtering.

### Case study

To demonstrate the power and application of direct Protein-SIF we applied it to a well-studied bacteria-animal symbiosis, the gutless marine oligochaete *Olavius algarvensis* (Fig. 4a). *O. algarvensis* lacks a digestive system. Instead, the worm relies on at least five bacterial symbionts under its cuticle for nutrition (Fig. 4b and c). The metabolism and physiology of these symbionts and their interactions with the host have been extensively studied using metagenomics, metaproteomics and physiological incubation experiments (15-17). Based on these previous studies, the carbon sources of the symbionts and host were thought to be as follows (Fig. 4c): Two sulfur oxidizers (γ1 & γ3) fix seawater derived inorganic carbon using the Calvin-Benson-Bassham cycle with a Form IA RubisCO enzyme (15). Two sulfate reducers (δ1 & δ4) consume the host’s organic waste products. For a spirochete, no data on metabolism and physiology was available. The host itself derives its carbon directly from the symbionts by digestion through endocytosis (18). Thus, the δ^13^C values of the γ- and δ-symbionts as well as the host were expected to be similar, since all carbon is derived from the initial carbon fixation by the γ-symbionts. In a previous study the δ^13^C value of complete worms was determined by IRMS to be -30.6‰ (16), so the expectation would be that all δ^13^C values are somewhere in the range of -25 to -35‰.

**Figure 4:**
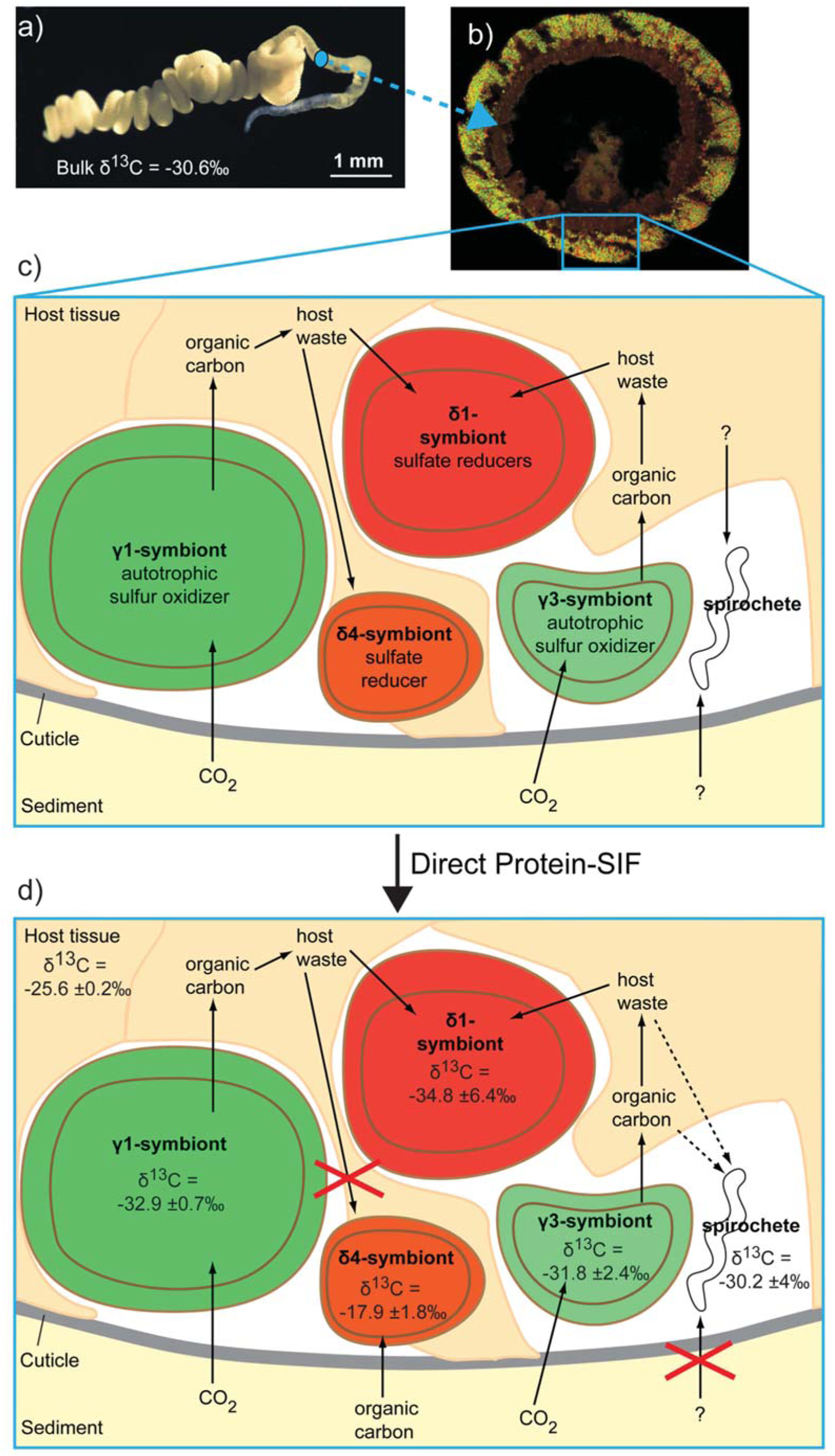
Testing the model of physiological interactions in the *Olavius algarvensis* symbiosis using direct Protein-SIF. a) Live *O. algarvensis* specimen. The δ^13^C value of bulk worms was determined by IRMS on six biological replicates in Kleiner et al. (2015) (16). b) Cross section through the worm. The bacterial symbionts right below the worm’s cuticle are stained with specific fluorescence *in situ* hybridization probes (γ-symbionts in green, δ-symbionts in red). c) Simplified model of carbon flow in the symbiosis based on previous metagenomic (17) and metaproteomic (19) studies. For the δ1- and δ4-symbionts the metaproteomic data suggested that these two symbionts are highly similar in terms of metabolism and physiology. d) Adjusted model of carbon flow based on carbon sources predicted using direct Protein-SIF derived δ^13^C values. Detailed Protein-SIF data in Table S5. The ‘±’ value for each δ^13^C value indicates the standard error.

We put this model of carbon flow in *O. algarvensis* to the test by applying direct Protein-SIF using a metaproteomic dataset obtained from multiple individual worms. To obtain sufficient peptides for measurement of δ^13^C values of all the symbionts, we combined 18 LC-MS/MS runs from a total of 14 individual worms. The δ^13^C values obtained by direct Protein-SIF of the host (-25.6 ‰), the two γ-symbionts (-32.9 and -31.8‰) and the δ1-symbiont (-34.8‰) were in the expected range (Figure 4d, Table S5). However, the δ4-symbiont had a much higher δ^13^C value (-17.9‰, Standard Error ±1.8‰) than the δ1-symbiont. This was unexpected because the two δ-symbionts appear to be almost identical in terms of expressed carbon uptake and catabolic pathways. Both are characterized by many abundantly expressed high affinity uptake transporters for sugars, amino acids, peptides and organic acids, as well as pathways for the use of host derived acetate and propionate (19). The direct Protein-SIF data now points towards a functional difference between the two δ-symbionts. Apparently, the δ4-symbiont derives part or all of its carbon from external sources that have a different carbon isotopic composition. We were also able to measure a sufficient number of peptides to estimate a δ^13^C value for the spirochetal symbiont, despite its low abundance. The spirochete δ^13^C value of -30‰ was in the range of symbiosis-internal carbon, suggesting that this symbiont uses a symbiosis-internal carbon source. Currently, we are not aware of any symbiosis-external carbon sources that have a δ^13^C value in the range of -30‰, however, we cannot exclude that the spirochete has access to a symbiosis external source with such a negative δ^13^C value. In summary, by applying direct Protein-SIF to the *O. algarvensis* symbiosis we derived new insights into carbon flow that were not suggested by any of the previous meta-omics based studies.

## Discussion

The developed direct Protein-SIF approach provides, for the first time, a means to directly and simultaneously access the SIFs (i.e. δ^13^C values) of many individual species in microbial communities. As little as 4 hours of LC-MS/MS time can be sufficient to estimate SIFs for the most abundant species in a sample. For this type of analysis, only very small sample amounts are needed; recent advances in sample preparation allow for the production of metaproteomic data of sufficient quality from as little as 1 mg of wet weight cell mass. If the determination of SIFs for lower abundant species is desired, more LC-MS/MS run-time can provide the required metaproteomic depth. For longer or additional LC-MS/MS runs proportionally larger amounts of sample are needed, for example, 2 mg for two 4-hour runs. Additionally, enrichment of specific cell populations by filtration or centrifugation methods can be used to obtain better metaproteomic coverage (19, 20).

We observed a range of differences between the δ^13^C values measured by IRMS and direct Protein-SIF when using both on single-species biomass (absolute deviation of the Protein-SIF δ^13^C values from the CF-EA-IRMS δ^13^C values was on average ±2.1‰). Part of the observed variation between the two methods is likely due to a lower accuracy of direct Protein-SIF using the highly-complex metaproteomic mass spectrometric data. However, there are at least two other factors that might contribute to this variation in the observed δ^13^C values. First, direct Protein-SIF measures the δ^13^C value of proteins, while bulk IRMS measures all cell components such as protein, lipids, DNA, and metabolites, providing a weighted average. It has been shown that the δ^13^C values of different cell components can differ. For example, lipids have a 1.6‰ lower δ^13^C value than protein in *Escherichia coli* (21) and the lipids of Calvin-Benson-Basham cycle autotrophs have around 6‰ lower δ^13^C values as compared to the δ^13^C values of the total biomass (5, 7). The possible difference between protein δ^13^C values and bulk organic matter δ^13^C values should thus be considered when interpreting direct Protein-SIF results. However, since protein usually makes up the majority of a cell in terms of mass, e.g. 55% of *E. coli* dry weight (BNID 104954)(22), protein δ^13^C values will be a good approximation of bulk δ^13^C values in most cases.

Second, the variation in the intensities of the peptide isotopic peaks used for direct Protein-SIF is mostly due to variation in the ratio of ^13^C to ^12^C, because ^13^C is, with a natural abundance of ~1.1%, the most abundant heavy stable isotope in the considered peptides. However, very large isotope fractionation of the three other elements (hydrogen, oxygen and nitrogen) in the peptides that we consider (sulfur containing peptides are excluded) could change the measured δ^13^C values by several per mille (‰). For example, the hydrogen isotope fractionation in photosynthate as compared to the water used by the photosynthetic organism is ε=-171‰ (5). The ^2^H/^1^H ratio in the Vienna Standard Mean Ocean Water (VSMOW) reference is 0.00015576, which means that the ^2^H/^1^H ratio of photosynthate is 0.000129 if ocean water is the hydrogen source i.e. a change in the fifth decimal of the fraction of heavy atoms. For comparison, carbon isotope fractionation influences the third and fourth decimal. As the ^13^C/^12^C ratio of Vienna Peedee Belemnite is 0.0111802, a δ^13^C value of -47.2‰ corresponds to a ^13^C/^12^C ratio of 0.01065. Carbon isotope fractionation by only -3‰ would already yield a ^13^C/^12^C ratio of 0.01115, a shift in the fraction of heavy atoms similar to that produced by the fractionating hydrogen isotopes by -171‰. Or, to put it differently a δ^2^H value of -171‰ would change the estimated δ^13^C value of -47.2‰ to -50.1‰ in direct Protein-SIF if the reference material used had a δ^2^H value similar to that of VSMOW (0 ‰). However, protein reference materials used for direct Protein-SIF will have a much more negative δ^2^H value as compared to VSMOW (e.g. the δ^2^H values of human hair range typically between -130 and -80‰ (23)) thus removing the most common hydrogen isotope fractionation effects when applying the reference material based offset correction to the direct Protein-SIF δ^13^C values. Therefore, while nitrogen, hydrogen and oxygen isotope fractionation will usually not have a major effect on direct Protein-SIF δ^13^C values, it should always be considered during interpretation of direct Protein-SIF results. Measurement of δ values for these three elements by IRMS bulk analyses of samples and reference material would provide insights into any major deviations that should be considered to achieve even better accuracy of the results.

The direct Protein-SIF approach provides us with a key ability that no other method can provide at the moment. By measuring δ^13^C values of individual species in microbial communities we can now make inferences about the food sources for these species as well as the metabolic pathways used for carbon assimilation. Ongoing technological development will further improve the accuracy and detection limits of the direct Protein-SIF approach in the future in at least two ways. First, the fractionation of isotopic species in the mass spectrometer could be reduced with specialized methods and potentially with improvements of the instrumentation. Current approaches for reducing isotope fractionation in Orbitrap mass spectrometers are very promising even though they are not yet useful for metaproteomics (12); these new approaches significantly reduce the number of MS^2^ spectra acquired and thus reduce the number of identified peptides. Second, mass spectrometers with increased resolving power at high scan rates will make it possible to separate the isotopologues of peptides based on which element provides the heavy isotope (hydrogen, nitrogen, carbon, oxygen). For example, the exchange of one ^14^N atom with one ^15^N atom in a peptide changes its mass by 0.99703489 Da, while the exchange of one ^12^C with one ^13^C changes the mass by 1.0033548378 Da. Once isotopic species can be resolved on a per-element basis, it should become feasible to not only determine carbon isotope ratios, but also isotope ratios for other elements in peptides such as nitrogen, oxygen and hydrogen.

## Online methods

### Sample preparation

The cultivation of the pure cultures and the creation of the mock communities are described in Kleiner et al. (2017) (14). For the case study, 14 *Olavius algarvensis* specimens were collected off the coast of Sant’ Andrea Bay, Elba, Italy (42°48’26’’N, 010°08’28’’E) in August 2015 from shallow-water (6 – 8 m water depth) sediments next to seagrass beds. Live worms were transported in native Elba sediment and seawater to the Max Planck Institute for Marine Microbiology in Bremen, Germany, and kept for one month in the dark and at Elba ambient temperature. Worms were then carefully removed from the sediment and frozen at -80°C until further processing.

The human hair used as a standard for correction of instrument isotope fractionation was obtained from the first author of this study (MK).

Peptide samples for proteomics were prepared and quantified as described by Kleiner et al. (2017) (14) following the filter-aided sample preparation (FASP) protocol described by Wisniewski et al. (24). The only modification for the *Olavius* samples as compared to the pure cultures and the mock communities was that no bead beating step was used. The bead beating step was used for the human hair reference material.

### 1D-LC-MS/MS

Samples were analyzed by 1D-LC-MS/MS as described in Kleiner et al. (2017) (14). One or two wash runs and one blank run were performed between samples to reduce carry over. For the 1D-LC-MS/MS runs of pure culture samples, 2 μg of peptide were loaded onto a 5 mm, 300 μm ID C18 Acclaim^®^ PepMap100 pre-column (Thermo Fisher Scientific) using an UltiMate™ 3000 RSLCnano Liquid Chromatograph (Thermo Fisher Scientific). After loading, the pre-column was switched in line with a 50 cm × 75 μm analytical EASY-Spray column packed with PepMap RSLC C18, 2 μm material (Thermo Fisher Scientific). For the 1D-LC-MS/MS runs of the *Olavius* samples, 0.8 to 4 μg of peptide were loaded onto a 2 cm, 75 μm ID C18 Acclaim^®^ PepMap 100 pre-column (Thermo Fisher Scientific) using an EASY-nLC 1000 Liquid Chromatograph (Thermo Fisher Scientific) set up in 2-column mode. The pre-column was also connected to a 50 cm × 75 μm analytical EASY-Spray column packed with PepMap RSLC C18, 2 μm material. In both cases the analytical column was connected via an Easy-Spray source to a Q Exactive Plus hybrid quadrupole-Orbitrap mass spectrometer (Thermo Fisher Scientific). Peptides were separated on the analytical column using 260 or 460 min gradients and mass spectra were acquired in the Orbitrap as described by Petersen et al. (2016)(25). For the mock communities, the existing 1D-LC-MS/MS data from Kleiner et al. (2017) (14) was used.

### Input data generation, algorithm and software development for Protein-SIF

#### Peptide identification

For peptide and protein identification of pure culture samples, protein sequence databases were created using the reference protein sequences for each species separately. The databases are available from the PRIDE repository (PXD006762). For the mock community samples, the database from the PRIDE repository (PXD006118) was used (14). For the Olavius samples, an existing *Olavius algarvensis* host and symbiont protein database from project PXD003626 (ftp://massive.ucsd.edu/MSV000079512/sequence/) was used. To this *Olavius* database we added additional symbiont protein sequences from recently sequenced metagenomes. The new *Olavius* database is available from the PRIDE repository (PXD007510). For the human hair standard, the human reference protein sequences from Uniprot (UP000005640) were used. CD-HIT was used to remove redundant sequences from the multimember databases using an identity threshold of 95% (26). The cRAP protein sequence database (http://www.thegpm.org/crap/) containing protein sequences of common laboratory contaminants was appended to each database. One important consideration for creating protein sequence databases for use with the Calis-p software is that Calis-p will calculate taxon/population specific δ^13^C values based on Accession number prefixes that indicate to which taxon a sequence belongs. The prefix should be separated from the accession number by an underscore (e.g. >TAX_00000). MS/MS spectra were searched against the databases using the Sequest HT node in Proteome Discoverer version 2.0.0.802 (Thermo Fisher Scientific) and peptide spectral matches were filtered using the Percolator node as described by Petersen et al. (2016)(25). The FidoCT node in Proteome Discoverer was used for protein inference and to restrict the protein-level FDR to below 5% (FidoCT q-value <0.05).

**Input files**: Examples for all input files are provided in PRIDE project PXD006762 alongside with the raw data. The LC-MS/MS produced raw files were converted into mzML format using MSConvertGUI via ProteoWizard (27) with the following options set: “Output format: mzML, Binary encoding precision:64-bit, Write index: checked, TPP compatibility: checked, Filter: Peak Picking, Algorithm: Vendor, MS Levels: 1 (The MS/MS scans are not needed for isotope pattern extraction). The peptide-spectrum match (PSM) files generated by Proteome Discoverer were exported in tab-delimited text format. The mzML files and the PSM files were used to extract isotopic patterns for all identified peptides.

#### Isotopic pattern extraction

The steps described here are carried out by the Calis-p software in a fully automated fashion upon provision of correctly formatted input files. The entries in the PSM files were excluded if they: (i) had any identified post-translational modifications (PTMs), (ii) contained “M” or “C” in the peptide sequence, (iii) had a peptide confidence score not equal to “High”, or (iv) had a PSM Ambiguity value equal to “Ambiguous”. The remaining entries were subjected to the isotopic pattern extraction from the mzML mass spectrum input files. In a first step, the m/z value of the monoisotopic peak (A) of each peptide was searched for in the full scans in the mass spectrum file in a defined retention time window (peptide scan start time ± 0.5). All peaks with the same m/z value in the retention time window were considered to be from the same peptide, because most peptides are analyzed multiple times in subsequent MS^1^ scans in the defined RT window size. In a second step, each MS^1^ scan of the same peptide was searched for the isotopic peaks of the peptide produced by replacement of single or multiple atoms in the peptide with heavier isotopes (i.e. A+1, A+2…). The isotopic peaks were defined as all the peaks following the monoisotopic peak in the full scan mass spectrum with m/z values being n*(1/charge) distance removed from the monoisotopic m/z value (n is the peak count i.e. A+1, A+2.). Since the number of isotopic peaks found for a peptide in different MS^1^ scans is variable due to decreasing signal intensity for the higher number isotopic peaks, we chose to report the peak intensities only for the isotopic peaks detected in the majority of scans. To clarify, if we found the A, A+1, A+2 peaks for five MS^1^ scans, A, A+1, A+2, A+3 peaks for three MS^1^ scans, A, A+1, A+2, A+3, A+4 for seven MS^1^ scans, and A, A+1, A+2, A+3, A+4, A+5 for three MS^1^ scans, we only reported peak intensities for A to A+4. Scans that did not have all isotopic peaks to be reported were excluded i.e. for the preceding example this would mean a scan that only had peaks A to A+3 would be excluded. To maximize carbon ratio calculation accuracy the intensities of each isotopic peak from all the included scans of the same peptide were summed up. The filtered PSM entries with the identified isotopic patterns and aggregated intensities were written to a isotopicPattern text file in a tab delimited format. Additional information needed for SIF computation such as peptide sequence, peptide sum formula, charge, and peptide identification quality score (Xcorr) was written into the isotopicPattern file. For examples of isotopic pattern files see PRIDE project PXD006762.

#### SIF computation

Peptide isotope patterns were filtered to remove exact duplicates (peptides associated with identical spectra). Also, the following default parameters were used: peptides associated with more than three proteins or that were identified with low confidence (Xcorr < 2.0), or had fewer than three isotopic peaks in their spectra, or had a total intensity (sum of all peaks) of less than 0.5 × 10^8^, were not considered further.

To calculate the δ^13^C value for peptides, we compared the experimentally derived isotope peak intensities with simulated isotope peak intensities. For the peptides remaining after filtering, the absolute intensity of each peak (mono-isotopic mass, A+1, A+2, etc.) was divided by the summed intensity of all peaks to obtain relative peak intensities. The ^13^C/^12^C ratio for each peptide was computed as follows: For ^13^C/^12^C ratios between 0 and 0.1 and fixed (natural) relative abundances of ^15^N, ^18^O, ^17^O and ^2^H, the theoretical spectrum (relative intensity of each peak) was computed using a Fast Fourier Transform modified after Rockwood et al. (1995)(11) based on the peptide’s molecular formula. In contrast to the approach used by Rockwood et al., we did not model the peaks as Gaussians to avoid the assumption that the peak for each mass had a Gaussian shape, as the experimental peaks clearly did not have a Gaussian shape. Instead, we simply computed the total intensity of each peak. Next, the theoretical peak intensities were corrected for the different number of peaks observed in the experimental spectrum and predicted for the theoretical spectrum. Finally, the goodness of fit between the experimental and theoretical spectrum was calculated as the sum of squares of the difference between observed intensity and predicted intensity for each peak. The goodness of fit was optimized (lowest sum of squares) by decreasing the step size of the ^13^C/^12^C value from 0.01 to 0.0001 in four iterations. For each species (population) in the sampled community (identified by an accession number prefix for the taxon), the average ^13^C/^12^C ratio was calculated as the intensity-weighted average of the ^13^C/^12^C ratio of each peptide attributed to that population. These data were also used to estimate the standard error and the δ^13^C for each population. To eliminate results from peptides suffering from interference from other peptides (due to overlapping spectra), only results with a sum of squares lower than 0.00005 or those results with a goodness of fit in the upper 33% of the data for that population were used in the calculation of the average δ^13^C values.

#### Correction of direct Protein-SIF values with reference standard

To correct for isotope fractionation introduced during sample processing and analysis, mostly by isotope fractionation in the mass spectrometer (12) (see also Results), we used human hair as a standard. δ^13^C values for the human hair standard were obtained both with direct Protein-SIF and CF-EA-IRMS (see below) and the offset between the two measurement methods was calculated. The offset value was then used to correct the direct Protein-SIF values obtained for the pure culture and for the community samples.

### Continuous Flow-Elemental Analysis-Isotope Ratio Mass Spectrometry (CF-EA-IRMS)

Stable isotope ratios of carbon and nitrogen of bacterial strains, human hair and culture media were determined (Tables S2 and S3) by CF-EA-IRMS. Frozen cell pellets of bacterial strains were dried at 105 °C and stored at 4 °C. Dried cell pellets as well as carbon and nitrogen sources used for cultivation were weighed (approx. 1 mg/sample) into tin capsules and stored in a desiccator. δ^13^C and δ^15^Ν values were measured using Continuous Flow-Elemental Analysis-Isotope Ratio Mass Spectrometry (CF-EA-IRMS) on a Delta V Plus (Thermo) mass spectrometer coupled to an ECS 4010 elemental analyzer (Costech). A ‘Zero Blank’ autosampler (Costech) was used for dropping the samples onto a quartz tube combustion column heated to 1000 °C. Simultaneously with sample dropping, an O_2_ pulse was injected to flash-combust the samples. Ultra-high purity helium was used as a carrier gas to transport the resulting gases through a water trap and a reduction furnace (600 °C) in which NOx species were reduced to N2. The generated N_2_ and CO_2_ were separated on a GC column. A Finnigan ConFlo III (Thermo) interface was used to deliver the gases to the ion source of the mass spectrometer. Peak areas for the isotopic species of sample CO_2_ and N_2_ were compared to those of reference gas peaks to determine the δ^13^C and δ^15^Ν values of the samples. International standards (Table S3) were used at the beginning and end of the measurement sequence to normalize the measurements to the internationally accepted delta (δ) scale with respect to V-PDB for δ^13^C and atmospheric N_2_ for δ^15^N. Additionally, caffeine (Sigma Aldrich, #C-0750) was included at the beginning and gelatin (Sigma Aldrich, #G-9382) after each 5^th^ sample as internal lab standards. The measurement uncertainty was equal or better than ± 0.15 ‰ for δ^13^C and δ^15^N values determined by IRMS.

### Data and software availability

The Calis-p software was implemented in Java and is freely available for download, use and modification at https://sourceforge.net/projects/calis-p/. The mass spectrometry proteomics data and the protein sequence databases have been deposited to the ProteomeXchange Consortium (28) via the PRIDE partner repository for the pure culture data with the dataset identifier PXD006762 [Reviewer access: log in at http://www.ebi.ac.uk/pride/archive/ with Username: and Password: aK0YbWbY], dataset identifier PXD006118 for the mock community data published previously in Kleiner et al. (2017) (14), and the Olavius algarvensis case study date with dataset identifier PXD007510 [Reviewer access: log in at http://www.ebi.ac.uk/pride/archive/ with Username: and Password: bSnKz1y6].

## Author contributions

M. K., conceived study, obtained and created bacterial stocks for pure culture and mock community experiments, designed experiments, generated mass spectrometric data, developed and tested the algorithms together with M.S. and X.D., wrote the paper with input from all co-authors; X. D., designed and implemented the algorithm to extract isotopic patterns from raw data, prepared downloadable software package and online wiki page on sourceforge, and revised manuscript; T. H. prepared samples for IRMS analyses; J.W., collected and processed *O. algarvensis* samples for metaproteomics; E.T., prepared proteomics samples, carried out experiments and operated mass spectrometer, revised the manuscript; B.M., provided IRMS analyses and revised manuscript; M.S., conceived study, developed and coded the algorithm for estimating δ^13^C values, revised manuscript.

## Acknowledgements

We are grateful to Jianwei Chen, Emmo Hamann, Marc Mussmann, Jessica Kozlowski, Sean Booth, Jessica Duong, Johanna Voordouw, Kenneth Sanderson, JoongJae Kim, Joenel Alcantara, Anupama P. Halmillawewa, Michael F. Hynes, and Heidi Gibson for donations of cultures for the mock communities’ Christine E. Sharp for help with bacterial stock collection, Jackie Zorz, Maryam Ataeian and Angela Kouris for help with proteomics sample preparation, Angela Kouris and Eric Miller for grammar and spellchecking of the manuscript, Christian Lott for providing the *O. algarvensis* image used in Figure 4, and Steve Taylor and his team at the University of Calgary Stable Isotope Laboratory for measurements using isotope ratio mass spectrometry.

Sequencing of new *Olavius algarvensis* metagenomes was conducted by the U.S. Department of Energy Joint Genome Institute and is supported by the Office of Science of the U.S. Department of Energy under Contract No. DE-AC02-05CH11231. This study was supported by the Campus Alberta Innovation Chair Program (M.S., X.D.), the Canadian Foundation for Innovation (M.S.), the NC State Chancellor’s Faculty Excellence Program Cluster on Microbiomes and Complex Microbial Communities (M.K.), a DAAD fellowship by the German Academic Exchange Service (T.H.) and the Natural Sciences and Engineering Research Council (NSERC) of Canada through a Banting fellowship (M.K.), an NSERC Undergraduate Student Research Award (E.T.) and a NSERC Discovery Grant to M.S. and B.M. We acknowledge the support of the International Microbiome Center as well as funding support from the Government of Canada through Genome Canada, the Government of Alberta through Genome Alberta, Genome Prairie’ Research Manitoba, and Genome Quebec. This research was undertaken thanks in part to funding from the Canada First Research Excellence Fund.

---

**Figure S1:**
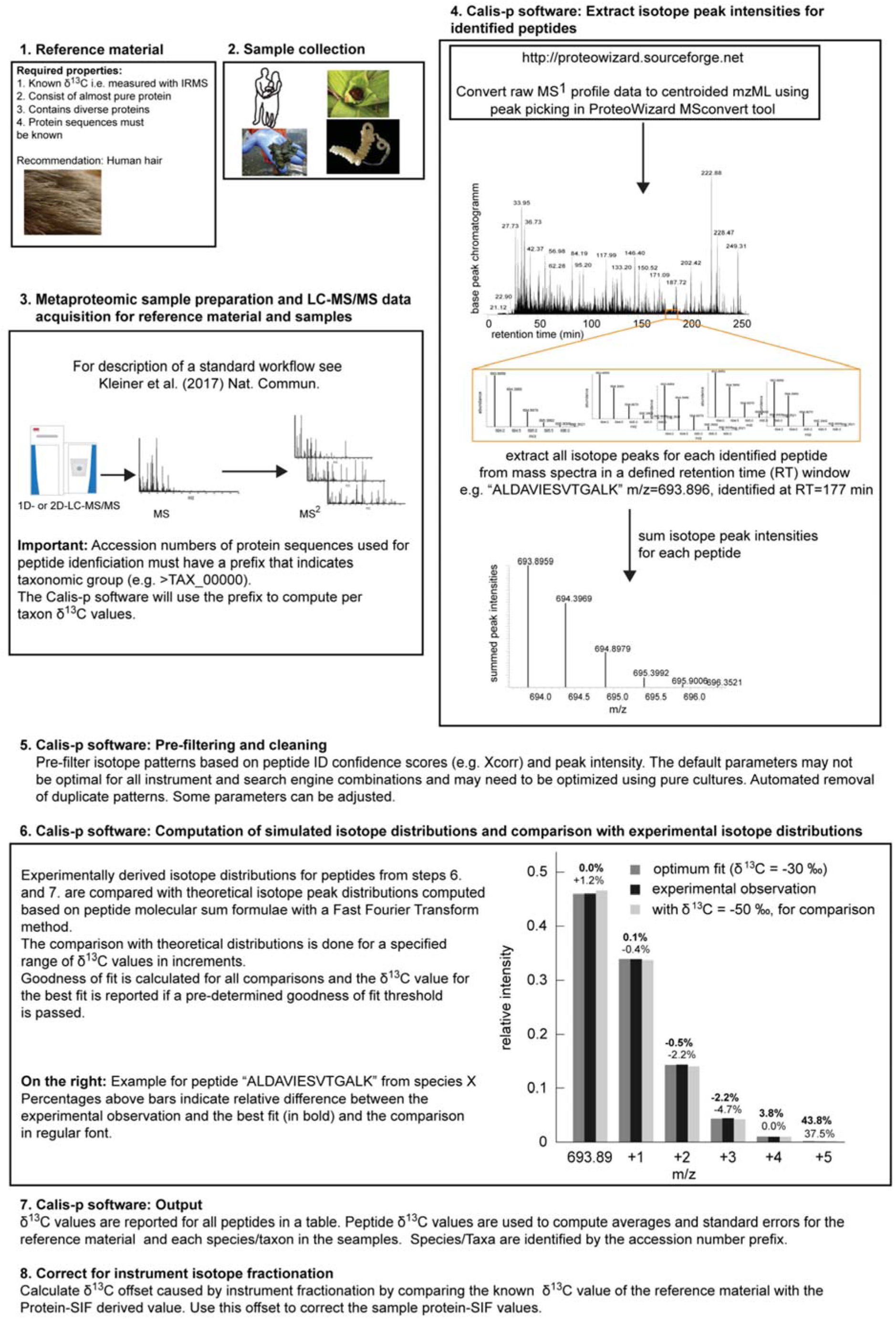
Direct Protein-SIF workflow with detailed explanations. Parts of step 3. were adapted from Kleiner et al. (14)

